# Flight trajectory modeling reveals species-specific obstacle avoidance policies in echolocating bats

**DOI:** 10.1101/2025.06.13.659477

**Authors:** Yu Teshima, Shoko Genda, Yota Aoki, Masahiro Fujisawa, Shizuko Hiryu, Keisuke Fujii

## Abstract

Echolocating bats routinely navigate complex wild environments with remarkable agility, yet it remains unclear whether their flight trajectories reflect reproducible internal control strategies or are merely the result of improvised reactive behavior. To address this question, we recorded flight paths and pulse emissions of *Rhinolophus nippon* and *Miniopterus fuliginosus* as they navigated seven obstacle-rich arenas in complete darkness. Using a machine learning model—a variational recurrent neural network (VRNN)—we show that bat flight is governed by a consistent internal policy. Trained on only part of each trajectory, our model accurately predicted future flight paths across arenas and individuals, preserving key features such as turning direction, obstacle-avoidance arcs, and velocity profiles. The model’s ability to faithfully reproduce species-specific strategies despite differences in sonar and flight morphology supports the existence of structured internal control in bat navigation. Our approach reveals that bat flight, while appearing improvisational, is in fact guided by internal policies that can be directly inferred from trajectory data, without relying on hand-crafted assumptions. Furthermore, the data-driven model developed here enables a priori testing of flight-path reorganization by virtually imposing environmental changes and analysing the resulting predicted trajectories, providing a powerful tool for biomimetic robotic design and for forecasting ecological responses to landscape transformation.

## Introduction

The sophisticated behavioral strategies exhibited by animals provide crucial insights into how different species adapt to their environments and ensure survival. Among these, echolocation in bats is widely recognized as a highly advanced sensorimotor integration strategy (1, 2). Echolocating bats emit pulses from their mouths or nostrils and analyze the returning echoes to perceive spatiotemporal structures in their surroundings. Despite the inherent physical constraints—including relatively low pulse-emission rate (3) and narrow beam directivity (4–7)—echolocating bats are capable of navigating complex environments with remarkable precision and flexibility (8–10). A deeper understanding of this capability not only holds ecological significance in elucidating the intellectual basis of bats but also offers valuable inspiration for engineering applications that aim to replicate such sensorimotor integration, including recent advances in deep reinforcement learning (11) and bio-inspired robotics (12–15).

Numerous studies have explored the mechanisms underlying echolocation-based navigation in bats. In particular, it has been hypothesized that flight paths shaped by active acoustic sensing are governed by specific internal policies (16), and many modeling efforts have aimed to formalize such strategies. Flight control for foraging (17–20), continuous prey pursuit (21), obstacle avoidance (22, 23), and light-dependent routing (24) have all been represented through analytical models, often in the context of bio-inspired robotics. Most of these approaches, however, rely on hand-crafted heuristics grounded in human intuition, which may obscure the flexibility, context sensitivity, and inter-individual variability characteristic of real bat behavior. Thus, whether bats indeed rely on reproducible internal control strategies, rather than improvisational reflexes, remains a fundamental open question.

To address this question, we examine the hypothesis that bat flight behavior follows predictable patterns indicative of underlying flight policies. If such patterns can be identified directly from behavioral data, it would suggest that bat navigation is organized around consistent sensorimotor strategies. To investigate this, we adopt a data-driven approach that extracts patterns from behavioral data and builds predictive models of flight using machine learning. We construct a set of obstacle-rich environments and record the flight trajectories and pulse emissions of two bat species, *Rhinolophus nippon* and *Miniopterus fuliginosus*, which differ markedly in both echolocation mode and flight dynamics. The collected data exhibit temporal structure, with flight trajectories and echolocation behavior varying dynamically in response to the bat’s state at each time step. These dynamics are captured using a variational recurrent neural network (VRNN) (25), a latent-variable time-series model previously shown to perform well in predicting structured movement (26–28). If the VRNN can accurately predict future trajectories from partial observations, this would provide quantitative support for the presence of internally structured, policy-like control in bat flight. We thus evaluate model accuracy by comparing predictions to actual trajectories, and interpret prediction errors as an indirect measure of how well internal control structure explains the observed behavior.

## Results

### Measurement set-up and behavioral overview

We measured the flight trajectories and pulse emission positions of two bat species—*Rhinolophus nippon* (*R. nippon*) and *Miniopterus fuliginosus* (*M. fuliginosus*)—in seven distinct obstacle environments (Env.1–7). These environments were constructed within an observation chamber equipped with 16 motion-capture cameras and 4 microphones, with chains suspended to form the obstacles (Fig. 1A). We recorded data from five individuals of *R. nippon* and four individuals of *M. fuliginosus*. Both species demonstrated skillful flight through the cluttered space without collisions (Fig. 1B).

**Fig. 1.**
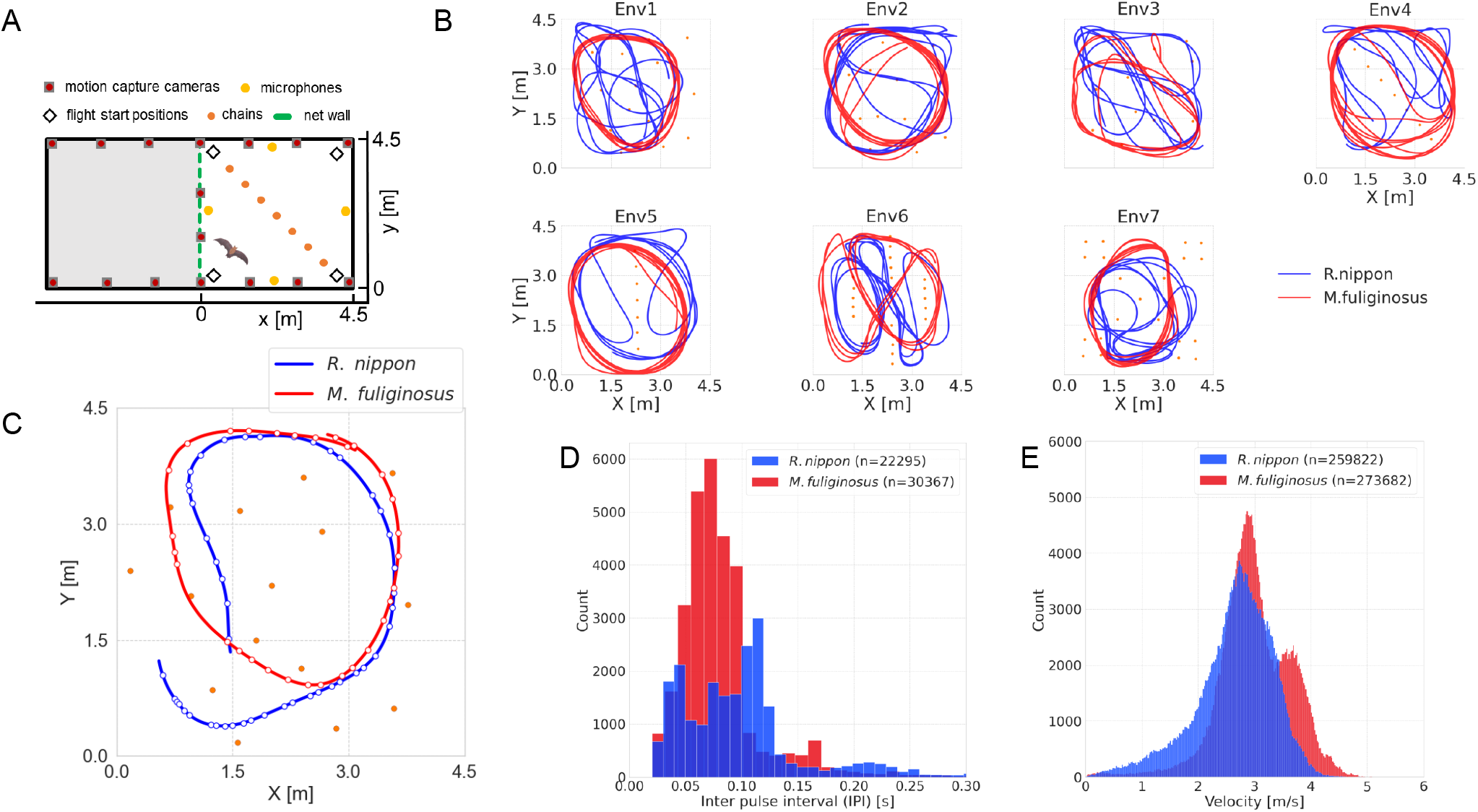
(A) Top view of the measurement system for flight trajectories and pulse emission timing of echolocation bats, in a 4.5 (x) m x 4.5 (y) m x 2.5 (z) m space separated from the observation room by a net. An obstacle space was constructed by hanging a 3.0 cm diameter chain as an obstacle from the ceiling. There were seven different obstacle spaces (Env 1 - 7). The bats started their flight from each of the four corners, and their flight trajectories were measured with infrared reflective markers attached to the bats’ backs, with motion capture located on the ceiling of the observation room, and pulse emission timing was measured with four microphones placed in the room. (B) Flight trajectories of a single individual of *R. nippon* and *M. fuliginosus*, measured over 12 s in each obstacle space; flight trajectories of both *R. nippon* and *M. fuliginosus* tended to converge to a certain trajectory over time, but the trajectory of *M. fuliginosus* were simpler than those of *R. nippon*. (C) Top view of the flight paths of *R. nippon* and *M. fuliginosus* over 4.0 s, and the locations of the emitted pulses (circular plots). (D) Histogram of inter-pulse interval (IPI) for *R. nippon* (n=22295) and *M. fuliginosus* (n=30367) for all trials; *M. fuliginosus* usually emits only a single pulse and therefore has a single peak in the histogram *R. nippon* emits double or triple pulses in addition to single pulses, so three peaks appear. The two species therefore differ not only in their flight tendencies but also in the frequency of information updates on the obstacle environment. (E) Histogram of velocity for *R. nippon* (n=259822) and *M. fuliginosus* (n=273682) for all trials.

To assess the frequency of spatial information acquisition, we calculated the inter-pulse interval (IPI) of echolocation. The average IPI was 1.1 *×* 10^2^ *±* 1.2 *×* 10^2^ ms for *R. nippon* and 90 *±* 90 ms for *M. fuliginosus*. Notably, in addition to single pulses, *R. nippon* frequently emitted double or triple pulses (Fig. 1C, D), indicating a distinct spatial sensing pattern compared to *M. fuliginosus*. In terms of flight speed, *R. nippon* flew at an average of 2.6 *±* 0.72 m/s, whereas *M. fuliginosus* exhibited a slightly higher average speed of 3.0 *±* 0.64 m/s (Fig. 1E, Table 1), often displaying relatively circular flight trajectories.

**Table 1.** The *L*_2_ predictions errors for the position and velocity of the *R. nippon* and *M. fuliginosus* using VRNN, and the predicted and measured velocities in each obstacle environment.

|  | *R. nippon* |  |  |  | *M. fuliginosus* |  |  |  |
| --- | --- | --- | --- | --- | --- | --- | --- | --- |
|  | Position error [m] | Velocity error [m/s] | Predicted velocity [m/s] | Measured velocity [m/s] | Position error [m] | Velocity error [m/s] | Predicted velocity [m/s] | Measured velocity [m/s] |
| All | $0.34 \pm 0.27$ | $0.88 \pm 0.53$ | $2.4 \pm 0.97$ | $2.6 \pm 0.76$ | $0.40 \pm 0.23$ | $0.97 \pm 0.52$ | $2.6 \pm 0.91$ | $3.0 \pm 0.61$ |
| Env1 | $0.29 \pm 0.16$ | $0.85 \pm 0.47$ | $2.4 \pm 0.91$ | $2.6 \pm 0.59$ | $0.34 \pm 0.18$ | $0.89 \pm 0.46$ | $2.4 \pm 0.73$ | $2.7 \pm 0.25$ |
| Env2 | $0.39 \pm 0.40$ | $0.89 \pm 0.63$ | $2.5 \pm 0.97$ | $2.7 \pm 0.88$ | $0.44 \pm 0.26$ | $1.0 \pm 0.56$ | $2.8 \pm 0.98$ | $3.3 \pm 0.70$ |
| Env3 | $0.29 \pm 0.19$ | $0.91 \pm 0.51$ | $2.6 \pm 1.0$ | $2.8 \pm 0.73$ | $0.42 \pm 0.23$ | $0.97 \pm 0.51$ | $2.7 \pm 0.89$ | $3.1 \pm 0.54$ |
| Env4 | $0.35 \pm 0.18$ | $0.82 \pm 0.46$ | $2.0 \pm 0.91$ | $2.1 \pm 0.61$ | $0.54 \pm 0.29$ | $1.2 \pm 0.62$ | $3.1 \pm 1.1$ | $3.7 \pm 0.86$ |
| Env5 | $0.43 \pm 0.24$ | $0.92 \pm 0.52$ | $2.5 \pm 0.96$ | $2.7 \pm 0.77$ | $0.45 \pm 0.20$ | $1.2 \pm 0.60$ | $3.2 \pm 0.97$ | $3.8 \pm 0.27$ |
| Env6 | $0.38 \pm 0.22$ | $0.84 \pm 0.43$ | $2.4 \pm 0.85$ | $2.5 \pm 0.52$ | $0.30 \pm 0.14$ | $0.80 \pm 0.45$ | $2.3 \pm 0.87$ | $2.5 \pm 0.60$ |
| Env7 | $0.49 \pm 0.35$ | $0.86 \pm 0.47$ | $2.1 \pm 1.1$ | $2.3 \pm 0.92$ | $0.34 \pm 0.18$ | $0.84 \pm 0.45$ | $2.4 \pm 0.86$ | $2.6 \pm 0.51$ |

### VRNN trajectory prediction

The recorded flight data were segmented into episodes of 8.0 seconds. The first 4.0 seconds of each episode were used as input to allow the model to perceive environmental information and update its internal state, while the subsequent 4.0 seconds were used for predicting flight velocities. The predicted velocities were integrated to obtain position coordinates, which were then compared to the actual measurements to evaluate prediction accuracy. Model performance was assessed using the *L*_2_-norm-based positional error, defined as the mean Euclidean distance between predicted and observed positions.

Predicted trajectories (red curves) aligned closely with the empirical flight paths (gray dashed lines) in nearly every test arena (Fig. 2, See Appendix, Fig. S1-2). The model faithfully reconstructed the intricate maneuvers imposed by obstacle geometry: in *R. nippon*, arenas Env2 and Env5, and in *M. fuliginosus*, arenas Env1 and Env4, it reproduced the overall path shape, turning sense, and obstacle-avoidance arcs. Minor, localized deviations appeared only in the most cluttered setting (Env6), yet the probabilistic envelopes generated by the VRNN still encompassed the observed tracks, leaving the global trajectory architecture virtually indistinguishable from the bats’ actual flights.

**Fig. 2.**
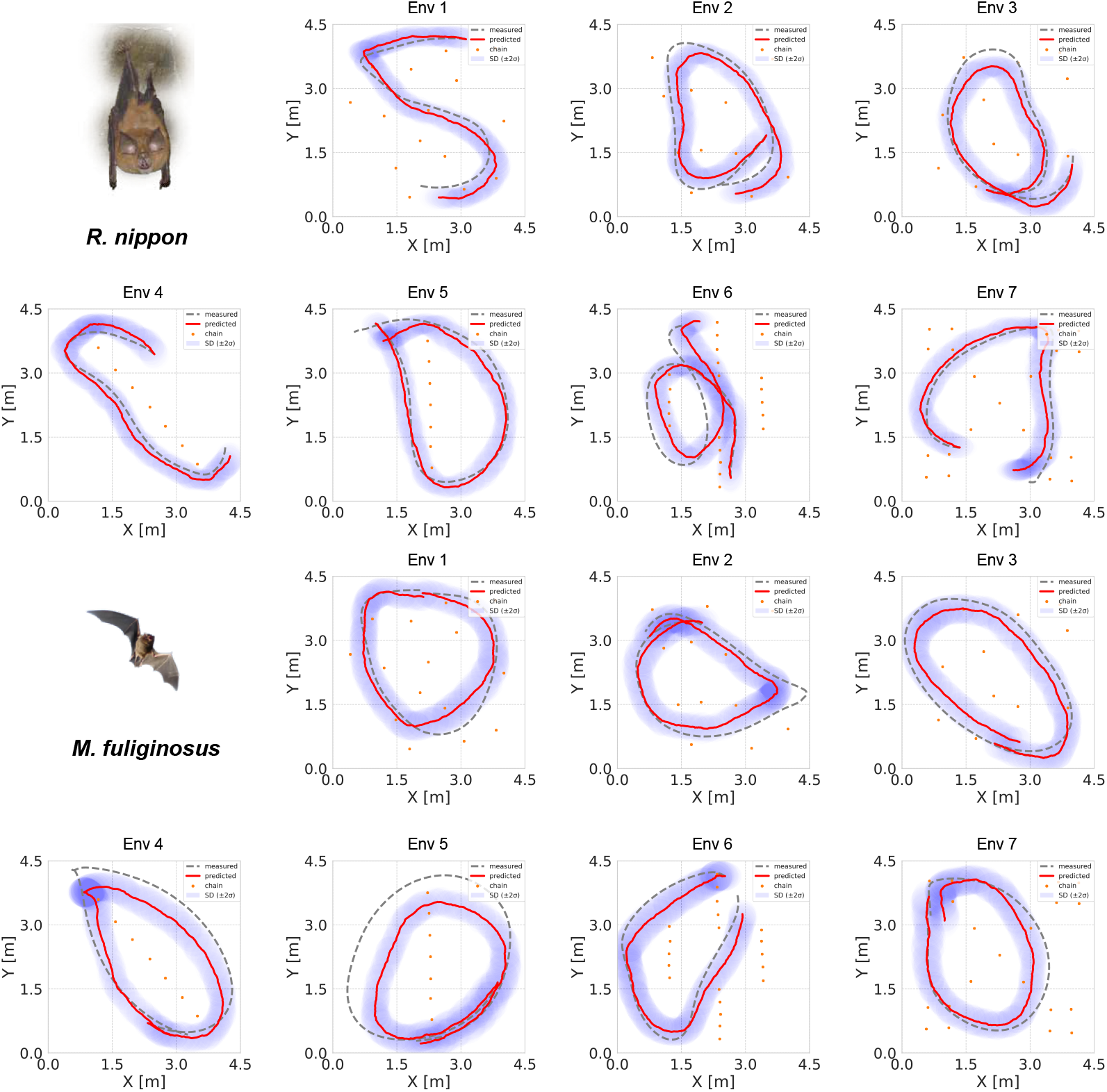
Top view of the 4.0 s measured flight trajectories (gray dotted lines) and sequentially predicted flight trajectories (mean: solid red lines and 2 times the standard deviation: blue) by VRNN for *R. nippon* and *M. fuliginosus* under four different obstacle environments (Env1 - 7). The input data to the model is 8.0 s, with the first 4.0 s being the burn-in period (29).

The average positional and velocity errors were 0.34 *±* 0.27 m and 0.88 *±* 0.53 m/s for *R. nippon*, and 0.40 *±* 0.23 m and 0.97 *±* 0.52 m/s for *M. fuliginosus*, respectively (Table 1). For both species, the model reproduced the general scale of flight speed accurately (actual vs. predicted: *R. nippon*: 2.6 m/s vs. 2.4 m/s; *M. fuliginosus*: 3.0 m/s vs. 2.6 m/s), with predicted values remaining close to the ground truth. In terms of environment-specific performance, *R. nippon* showed the smallest positional error in Env1 and Env3 (both 0.29 m), likely due to the relatively regular flight paths observed in these settings (Fig. 3). In contrast, errors were higher in Env7 (0.49 m) and Env5 (0.43 m), possibly due to the increased complexity and localized fluctuations of flight trajectories. For *M. fuliginosus*, good accuracy was achieved in Env6 (0.30 m) and Env1 (0.34 m), whereas the largest positional error occurred in Env4 (0.54 m), where predicted velocities also deviated substantially from the actual values (predicted: 3.1 m/s; actual: 3.7 m/s).

**Fig. 3.**
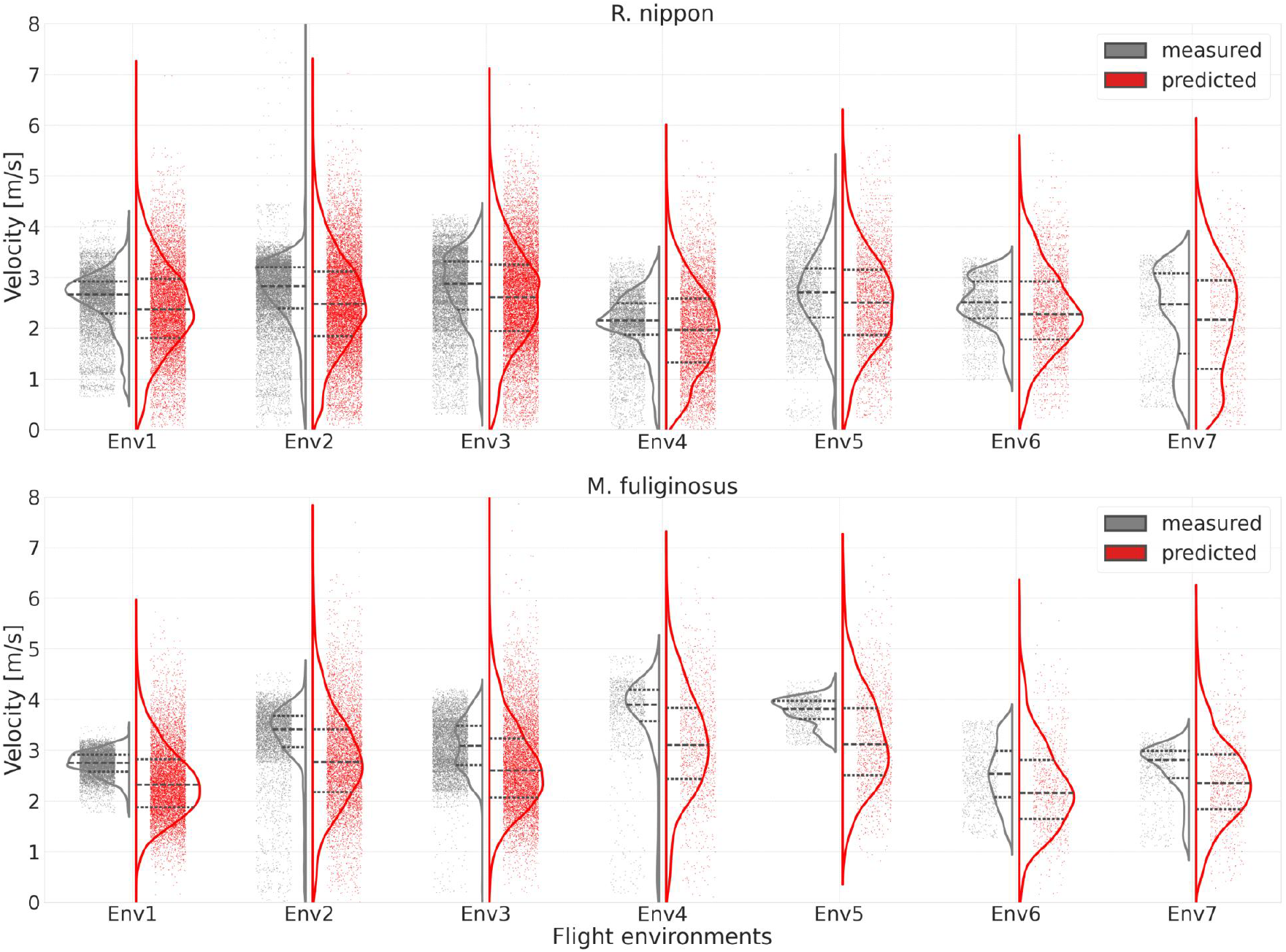
Measurement velocity (gray dots) and predicted velocity (red dots) of *R. nippon* and *M. fuliginosus* in each obstacle environment (Env1-Env7) and violin plots for each.

## Discussion

We investigated whether echolocating bats follow internally consistent flight policies by training the VRNN on flight trajectories recorded under diverse environmental conditions and environmental information calculated at pulse emission times. The model successfully reproduced species-specific flight behaviors—including turning direction, obstacle-avoidance arcs, and velocity profiles—for two bat species with distinct echolocation strategies. These results suggest that bat flight is not governed by random reactive control but instead by reproducible patterns, pointing to the presence of shared internal strategies across individuals and environments. While previous studies have inferred latent structure from recursive trajectories or implicated spatial memory cells such as place cells (30, 31), our data-driven approach directly extracted behavioral regularities from the trajectory data itself. Using machine learning as a pattern-recognition tool, we showed that seemingly improvisational flight paths can be predicted based on an underlying, consistent control policy.

The high-precision VRNN developed in this study not only corroborates the presence of internally consistent flight policies but also quantitatively predicts how bat trajectories reorganize under novel environmental conditions, including artificially designed scenarios. When trained on flight data from randomly arranged obstacle arenas (ENV 1–3) and tested on regularly arrayed arenas (ENV 4–7), the model maintained substantial predictive accuracy. The mean positional and velocity errors were 0.43 *±* 0.28 m and 0.94 *±* 0.57 m/s for *R. nippon*, and 0.58 *±* 0.35 m and 1.1 *±* 0.64 m/s for *M. fuliginosus*, respectively (See Appendix, Fig. S3–S4). This versatility distinguishes our data-driven framework from heuristic rule-based simulators. It can extrapolate how bats actually adjust their flight under unfamiliar conditions, offering new potential for biomimetic applications that exploit their adaptive control strategies. Quantifying the ecological impacts of wind-power expansion and artificial lighting on bat populations has become a pressing research priority (32, 33). Our framework, by learning from current data and predicting flight paths under future landscape changes, offers an evidence-based tool for evaluating the ecological consequences of human development. Such predictions can inform mitigation measures and benefit not only bats but also other species affected by habitat transformation.

A key strength of the VRNN framework is its ability to produce approximate predictive distributions over future velocity vectors, allowing us to quantify uncertainty through sampling. When the training data provides sufficient constraints, the predictive distribution becomes sharp, reflecting low aleatoric uncertainty. By contrast, when the model encounters conditions outside its training distribution or faces latent influences not captured in the input data, the distribution becomes broader—an indication of epistemic uncertainty. This distinction enables us to assess the reliability of individual predictions and to identify environments or species where the model’s confidence breaks down, highlighting gaps in data coverage or representational capacity.

In addition, combining our prediction model with a forward model of the bat’s auditory system (34–36) could simulate how acoustic inputs guide three-dimensional trajectories. Such a hybrid approach would not only deepen our understanding of internal flight policies but also represent a broader step toward mechanistic explanations of sensory-guided navigation in echolocating animals. Exploring this direction further constitutes an important avenue for future research.

While our model supports the existence of structured flight policies, it does not reveal what those policies actually entail. Deep learning models like the VRNN are powerful for prediction but often lack interpretability. At present, it remains difficult to determine which features in the input data most influence the model’s output, or how they contribute to the inferred control strategies. This limitation underscores a broader challenge in linking observed behavior to internal computation and points to the need for more interpretable modeling approaches. A promising way to tackle this challenge is to incorporate interpretable statistical frameworks that emphasize explanatory insight. Given that our study has established the existence of bat flight policies, hierarchical Bayesian models that capture species- and environment-level variation, or models that incorporate morphological traits such as body size, can simultaneously furnish interpretable structure and principled uncertainty quantification in modeling flight behavior.

## Conclusion

Our findings demonstrate that the seemingly improvisational flights of *R. nippon* and *M. fuliginosus* are orchestrated by stable internal control policies. The VRNN, trained on only the first half of each trajectory, reproduced the remainder with sub-meter accuracy across seven cluttered arenas. The model also separated aleatoric and epistemic uncertainty, revealing where predictions remain robust and where data gaps persist. This data-driven framework not only verifies the existence of reproducible strategies in echolocating bats but also provides a practical tool for forecasting behavioral responses to novel ecological threats, such as expanding wind-turbine arrays and increasing artificial lighting. Looking ahead, coupling learned control policies with models of biosonar reception represents a promising path toward interpretable frameworks that bridge neural implementation, behavioral flexibility, and ecological resilience in sensory-guided navigation.

## Materials and Methods

### Behavioral Experiments

#### Study Species

Two species of adult bats were included in this experiment. One is *Miniopterus fuliginosus* (*M. fuliginosus*) (one male and three females) and the other is *Rhinolophus nippon* (*R. nippon*) (five males). *M. fuliginosus* emits frequency-modulated (FM) pulses, and *R. nippon* emits a combination of constant-frequency (CF) pulses with FM pulses before and after the CF pulse. The flight characteristics of bats can be classified according to wing loading and aspect ratio. Wing loading is weight divided by wing area; the greater the weight, the faster the bat can fly. *M. fuliginosus* has high wing loading and aspect ratio, making it good for long-range fast flights, but not so good for sharp turns. *R. nippon* has low wing loading and aspect ratio, allowing it to change direction with a small turning radius and fly at low speeds. In a cave in Sabae City, Fukui Prefecture, Japan, three *M. fuliginosus* were captured on September 22, 2021, and one on September 14, 2020. One *R. nippon* was captured in the same cave on May 10, 2022, and four on May 16, 2023. All bats were housed in a dedicated colony room (4 m (L) x 6 m (W) x 2 m (H)) at Doshisha University, Kyoto. The colony room temperature was maintained at approximately 21° C and the humidity at 80% or higher. The bats could fly freely inside the colony room, consuming mealworms and water without restriction. The lighting in the room was also controlled to provide a 12 hour light / dark cycle. All necessary permits to capture and house the bats were obtained and the experiment was properly conducted under Japanese animal experimentation laws. Prior approval was obtained from the Animal Experiment Committee of Doshisha University.

#### Experimental Setup and Procedure

Behavioral experiments were conducted in seven different spaces with several obstacles (0.03 m diameter, plastic chains) placed in different positions in the observation room (4.5 m (L) x 4.5 m (W) x 2.5 m (H)) (Fig. 4A). The obstacles in Envs. 1, 2, and 3 were randomly placed, both in position and in number. In contrast, Envs 4, 5, 6, and 7 have regularly arranged obstacles (Fig. 4B). Bats started their flight at four locations at a height of 1.5 m in the four corners of the space. In each obstacle environment, bats flew four times, one trial from each of the four flight start positions. Bat flights were tracked using a 16-camera motion-capture system (Naturalpoint, Inc. DBA OptiTrack., Motive Tracker 3.0.1, USA) until the bats perched on a wall or net and stopped flying, and 3D flight trajectories were recorded at 100 frames / s. Flights in which the bat took off and landed within eight seconds were excluded from analysis. The motion capture camera was placed at a height of 2.4 m. For tracking, a 7.8 mm diameter reflector was mounted on the bat’s back. To eliminate as much as possible the influence of the experimenter on the release of bats from the starting point, flights immediately after release were excluded. Four microphones (Knowles, Fg-23329-D65) were placed at a height of 1.3 m around the observation space and recorded bat pulses emitted in flight at a sampling frequency of 500 kHz (National Instruments, PXIe-1073).

**Fig. 4.**
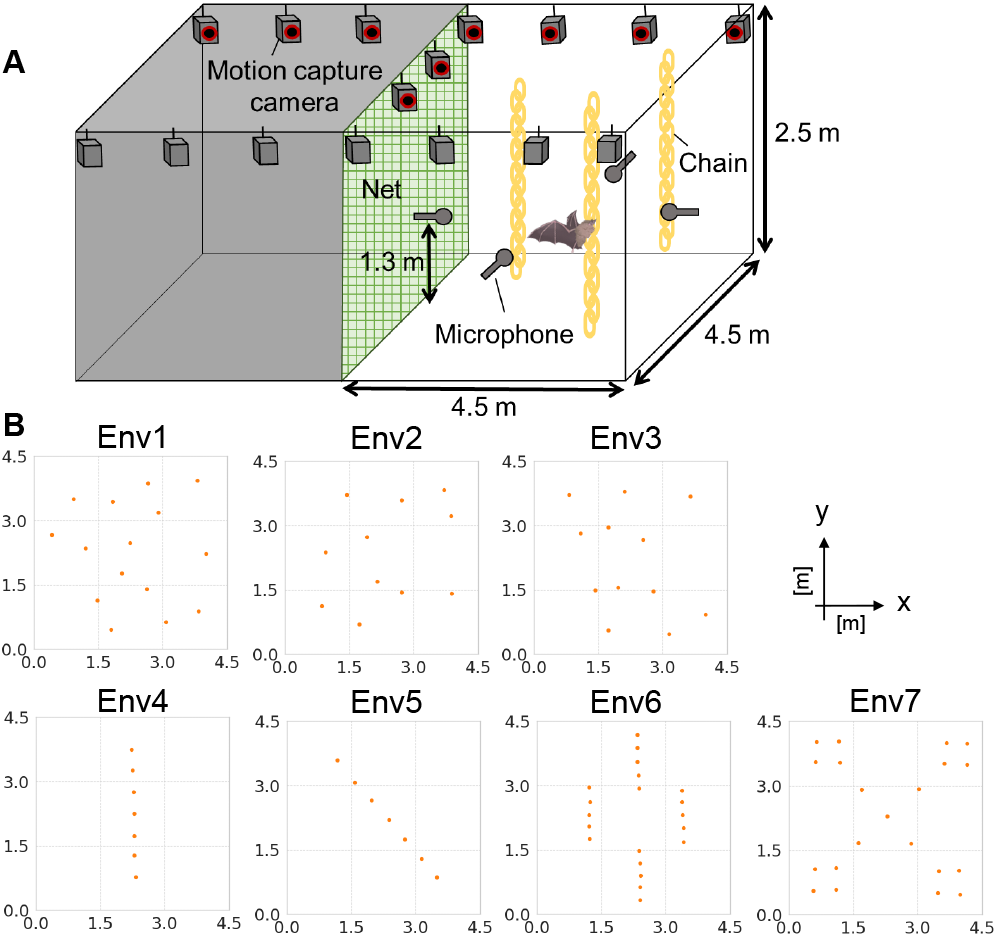
(A) Measurement system for flight trajectories and pulse timing of echolocation bats. (B) Top view of seven types of obstacle space (Envs 1 - 7).

#### Video and Audio Analysis

The bat flight trajectories measured by the motion capture system were smoothed and saved to CSV files using the smoothing function in the Motive Tracker application. Pulse emission times were calculated in MATLAB by applying a band-pass filter (65–70 kHz for *R. nippon*; 30–85 kHz for *M. fuliginosus*), converting the filtered signal to a spectrogram via fast Fourier transform (FFT), and locating the peak sound-pressure of each pulse. The FFT window length was 16384 points (frequency resolution: 30.5 Hz), and the shift length was 8192 points (time resolution: 16.4 ms). The time was down-sampled to fs = 100 Hz to synchronize with the flight trajectory and then saved to CSV files.

### Modeling flight policy by Neural Networks

#### Dataset

The dataset for the obstacle avoidance model was created from flight trajectories obtained in behavioral experiments. The *R. nippon* dataset consisted of 214 training data episodes (validation data: 20 episodes) and 56 episodes of test data, while the *M. fuliginosus* dataset consisted of 246 episodes of training data (validation data: 23 episodes) and 64 test data episodes (8.0 s / 1 episode). The dataset consists of the bat’s “action” and the “pulse states” of the bat when the action is taken in the two dimensions of the x-y plane. The “action” consists of the position coordinates (*x, y*), the flight speeds ((*V*_*x*_, *V*_*y*_)), the flight turn angles (*θ*), and the timing of the pulses (0 (not emit the pulse) or 1 (emit the pulse)). The “pulse states” is the environmental information at the time the bat emitted the pulse. The reason why position coordinates are used as input here is that bats have place cells (37) and grid cells (38). To obtain the “pulse state”, we first placed a total of 251-line segments, 5.0 m long, at 0.32° intervals over a directional range of approximately ±40° of the *M. fuliginosus* pulse (39) in the direction of flight from the location where the bat emitted the pulse (See Appendix, Fig. S3(A)). This interval width was set to allow the detection of obstacles (0.03 m in diameter, plastic chains) located 5.0 m away from the bat (40). The distance between the 251 line segments and the obstacle or wall was a normalized distance of 0.0 - 1.0, with 5.0 m as 1.0. Significant figures were set at three decimal places. For line segments that did not intersect an obstacle, the distance was set to 2.0. Since the directivity of *R. nippon*’s pulse is approximately *±* 20° (6), the values for line segments in the + 20° to + 40° and - 20° to - 40° ranges were always set to 2.0 (See Appendix, Fig. S3(B)). Finally, the data set consisted of 800 frames per episode and 8.0 seconds per episode. For the training data to build the flight policy model under obstacle environments, 80% of the data measured in seven obstacle environments (Env 1-7) were randomly selected, and 10% of them were used as validation data. The remaining 20% was used as test data.

#### Structure of Neural Networks

The VRNN architecture was based on previous studies (28). The input comprised 257 dimensions (“action” + “pulse states”), with the deterministic function of the RNN, estimated using the Gated Recurrent Unit (GRU) (41). The hidden state dimension was set to 100 dimensions, and the hidden layer was configured with 64 dimensions. The model parameters were initialized using a normal distribution using the Xavier method (42). The termination criterion for model training was based on two conditions: first, when the loss observed in the validation dataset decreased to less than 1.0 *×* 10^*−*4^ compared to the loss update ratio from the preceding epoch, and second, when the training process extended to 500 epochs. However, all training was completed before the epoch reached 500.

## Data Availability

All data supporting the conclusions of this study were deposited in an open-access repository and are accessible under the DOI doi:10.6084/m9.figshare.29209493. The analysis scripts and custom code are available on GitHub (https://github.com/tsmyu/PO-MC-DHVRNN (tag: v1.1)) and have been archived on Zenodo with versioned DOI doi:10.5281/zenodo.15638675. Together, these resources contain every file and detailed instruction required to reproduce our results. Further information is available from the corresponding author upon reasonable request.

## ACKNOWLEDGEMENTS

This work was supported by JSPS KAKENHI Grant Numbers 21H05300, 21H05295, 22H01503, 25H00745, 25K21286 and JST, ACT-X Grant Number JPM-JAX23CI.

## Supplementary Note 1: Supplementary Information

**Fig. S1.**
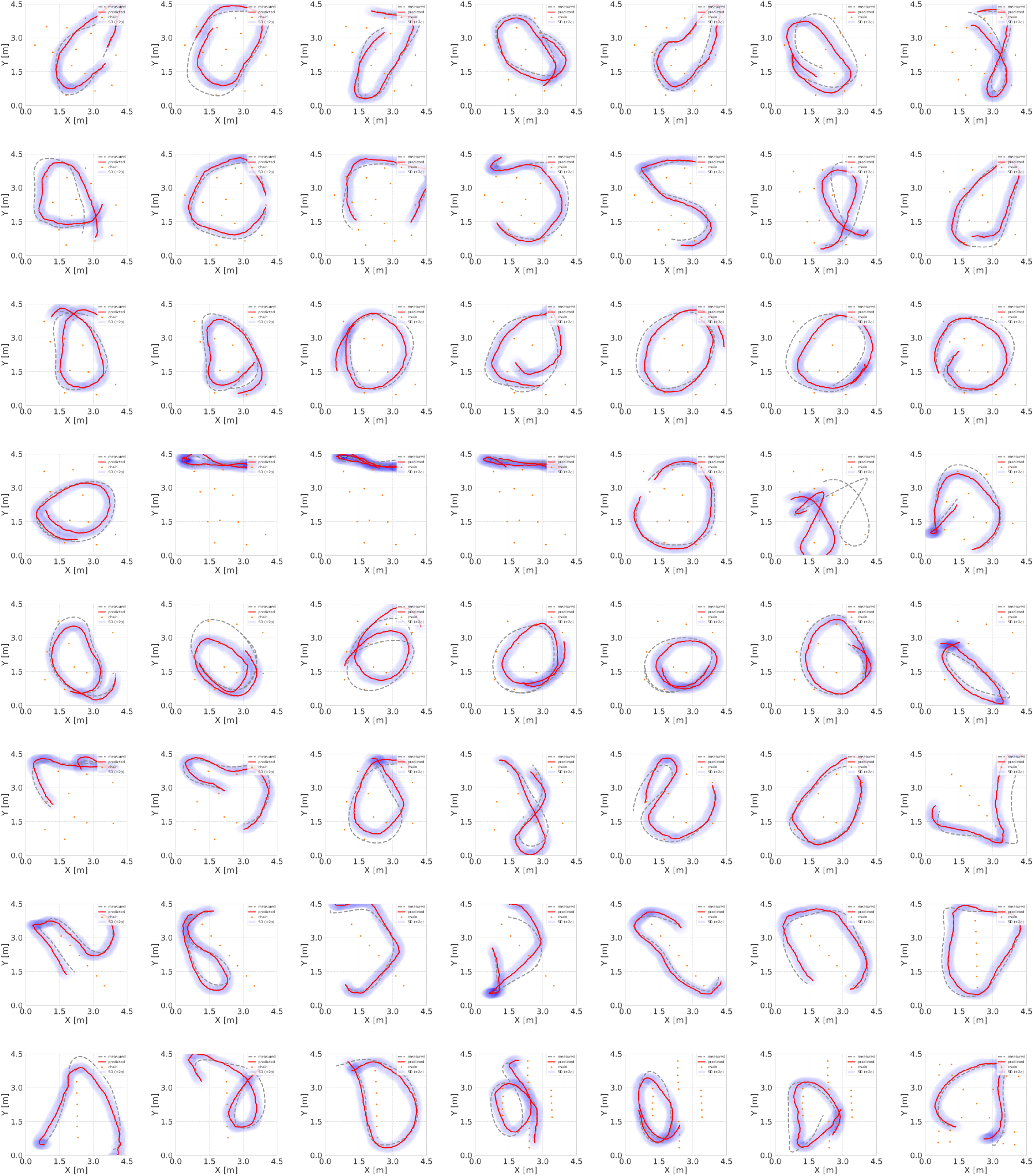
Top view of the measured flight path of *R. nippon* over 4.0 s (gray dotted line) alongside the trajectory predicted by the VRNN (training data: *R. nippon*) (red solid line). The input data to the model is 8.0 s, with the first 4.0 s being the burn-in period.

**Fig. S2.**
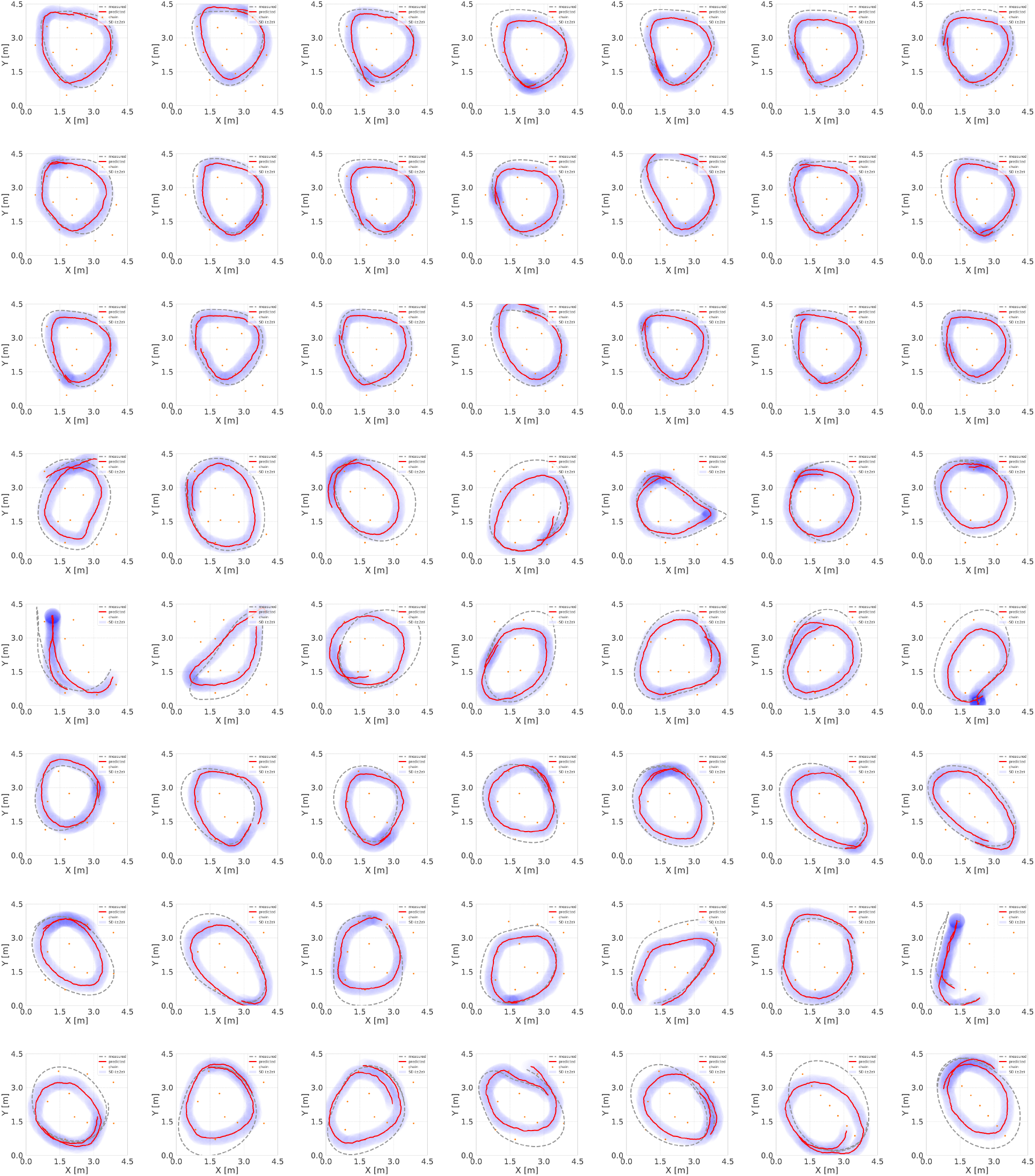
Top view of the measured flight path of *M. fuliginosus* over 4.0 s (gray dotted line) alongside the trajectory predicted by the VRNN(training data: *M. fuliginosus*) (red solid line). The input data to the model is 8.0 s, with the first 4.0 s being the burn-in period.

**Fig. S3.**
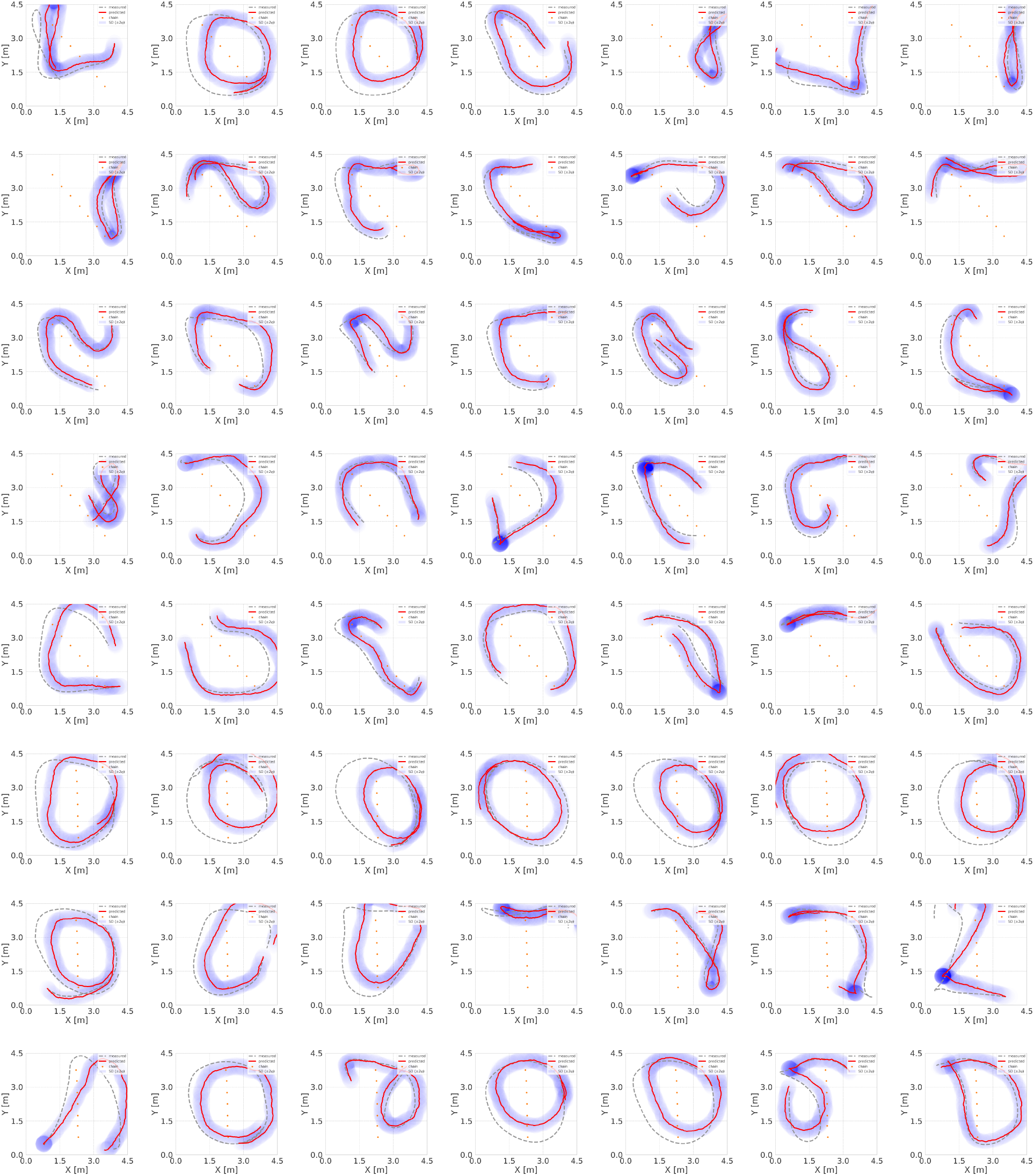
Top view of the measured flight path of *R. nippon* over 4.0 s (gray dotted line) alongside the trajectory predicted by the VRNN (red solid line). Data collected under environmental conditions ENV 1–3 were used for training, and those from ENV 4–7 for testing. The input data to the model is 8.0 s, with the first 4.0 s being the burn-in period.

**Fig. S4.**
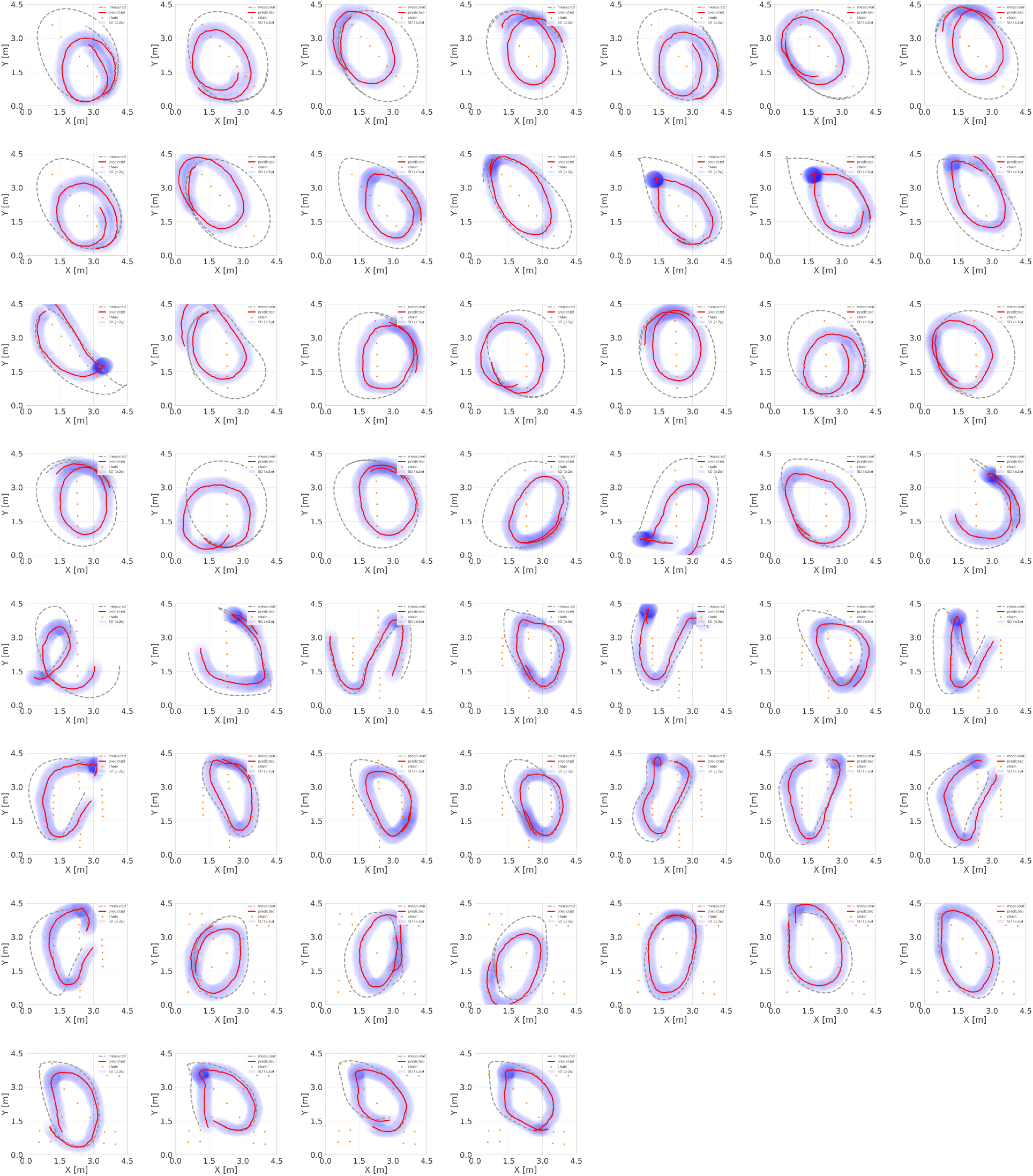
Top view of the measured flight path of *M. fuliginosus* over 4.0 s (gray dotted line) alongside the trajectory predicted by the VRNN (red solid line). Data collected under environmental conditions ENV 1–3 were used for training, and those from ENV 4–7 for testing. The input data to the model is 8.0 s, with the first 4.0 s being the burn-in period.

**Fig. S5.**
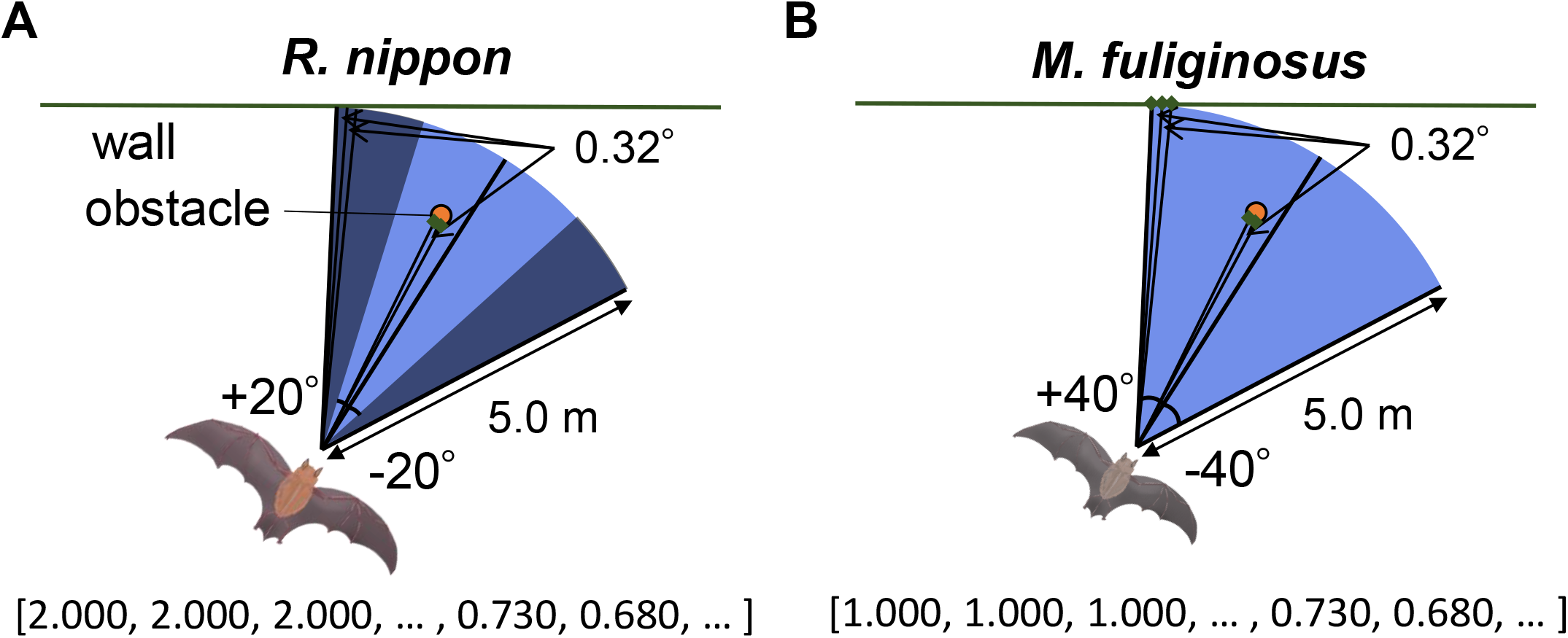
The following example illustrates the “pulse states” of *R. nippon* (A) and *M. fuliginosus* (B). The directionality of *M. fuliginosus* is ±40°, whereas that of *R. nippon* is ±20°. Consequently, the outer 20°of the *R. nippon* “pulse states” is consistently obstacle-free.

## Notes

### Competing Interest Statement

The authors have declared no competing interest.

### Summary of Updates

The tag version on the GitHub page has been changed, and the Grant number has been corrected. And the Subject Area is altered.

